# Probabilistic dietary exposure modeling and health risk assessment of heavy metals via the fodder–cattle–human continuum in Bangladesh

**DOI:** 10.64898/2026.05.30.728926

**Authors:** Shah Md. Iqbal, Md. Rakib Hasan, Kazi Rafiq, Anan Binte Zaman, Fardina Sultana Sumi, Md. Shafiqul Islam, Muhammad Tofazzal Hossain, A K M Anisur Rahman

## Abstract

Dietary exposure to heavy metals (HMs) via animal-source foods is a critical environmental health pathway. In rapidly industrializing Bangladesh, contamination of the bovine food chain from agricultural feeds and industrial emissions poses an unquantified public health burden. This study evaluated exposure pathways, spatial distribution, mass-transfer dynamics, and health risks of six HMs (Cr, Cu, Cd, Pb, As, and Hg) across the fodder–cattle–human continuum. Samples of beef (n = 76), raw milk (n = 76), commercial cattle feed (n = 40), and fodder (n = 88) were collected from eight sites across industrial and non-industrial zones in Bangladesh and analysed by atomic absorption spectroscopy. Probabilistic Monte Carlo simulations (10,000 iterations) quantified estimated daily intake, target hazard quotients (THQ), cumulative hazard index (HI), and lifetime carcinogenic risk (CR) for adult and pediatric receptors. Copper (Cu) was the dominant contaminant across all matrices, peaking in beef (103.89 ± 15.87 mg/kg) and milk (13.67 ± 1.53 mg/L). Spatial analysis revealed distinct contamination profiles: Pb burden peaked in industrial zones while Cr was elevated in non-industrial sectors. Monte Carlo modelling identified commercial feed as the most efficient transfer vector into beef. Pediatric THQ for Cu significantly exceeded the safety threshold (THQ > 1), and upper-bound lifetime carcinogenic risk from As approached the critical USEPA 10⁻⁴ regulatory ceiling. These findings demonstrate that industrial and agricultural externalities efficiently contaminate the bovine food supply chain in Bangladesh, with copper and arsenic representing the most critical non-carcinogenic and carcinogenic dietary hazards, respectively. Children are disproportionately vulnerable due to lower body weight. The results underscore the need for targeted upstream interventions in commercial feed production and provide evidence to support feed-quality regulation and environmental monitoring in rapidly industrializing settings.

## 1. Introduction

Heavy metals (HMs) are conventionally defined as metallic elements with a specific gravity exceeding 5 g/cm³ or atomic weights ranging from 63.546 to 200.590 g/mol. Even at trace concentrations these elements are toxic to living organisms [1–3] and persist in the environment owing to their non-biodegradable and non-thermodegradable properties [4]. Based on biological function, HMs are broadly classified as essential or non-essential. Essential trace elements—including cobalt (Co), chromium (Cr), copper (Cu), iron (Fe), manganese (Mn), molybdenum (Mo), nickel (Ni), selenium (Se), and zinc (Zn)—are indispensable for normal physiological processes when present at appropriate concentrations [5]. By contrast, non-essential metals such as arsenic (As), cadmium (Cd), mercury (Hg), and lead (Pb) serve no known biological function and pose significant toxicological risks even at low exposure levels.

The degree of HM toxicity varies considerably among elements and exposure levels. The Food and Agriculture Organization (FAO, 1999) ranks Cd as extremely toxic, followed by Pb (hazardous), As (moderately toxic), and Cr (somewhat toxic). Individual metals exert distinct biological effects: Cu promotes microbial growth at sub-inhibitory concentrations but functions as a potent antimicrobial agent at elevated levels [6], while Cd exerts cytotoxic effects even in trace amounts [7]. The World Health Organization lists As, Pb, Hg, and Cd among the ten most hazardous chemical contaminants in the human food supply, with well-documented health consequences including immunosuppression, carcinogenesis, hyperkeratosis, and multi-organ failure [8–10]. In livestock, acute exposure can be rapidly lethal; cattle have died within 24 hours of ingesting supralethal concentrations of Pb [11]. Anthropogenic activities are the primary driver of environmental HM dissemination. Historically, metals such as Hg, Cu, silver (Ag), and As have been deliberately introduced into agriculture and medicine as biocides, therapeutic agents, feed additives, and growth promoters [12,13]. In contemporary livestock production, copper sulfate is routinely incorporated into commercial feeds as a growth promoter and antifungal agent and is widely applied in dairy milking facilities as a foot-bath disinfectant [14]. Arsenical compounds have similarly been added to animal feeds to improve growth rate and feed conversion efficiency [15]. Beyond agriculture, HMs are released into the environment through industrial operations, including tanneries, battery recycling, metallurgical processing, and fossil fuel combustion [16,17].

Once released, HMs accumulate in soils and water bodies via the land application of sewage sludge, discharge of industrial effluents, and atmospheric deposition. Owing to their non-biodegradable nature, these elements persist in terrestrial and aquatic ecosystems for extended periods and are progressively absorbed by plants, entering the food chain [18,19]. Livestock are exposed primarily through ingestion of contaminated pasture grass or fodder irrigated with polluted wastewater, the most significant pathway for HM entry into the bovine food chain [20]. Dietary uptake results in bioaccumulation of Cd, Pb, Cu, Zn, Fe, Cr, and Mn in edible tissues, particularly muscle (beef) and milk [21]. Human dietary exposure to these metals occurs predominantly through consumption of contaminated animal products and, to a lesser extent, through inhalation, dermal contact, and drinking water [22].

Given the well-established toxicity of HMs at low doses, contamination of meat and milk represents a significant and growing public health concern [23,24]. Effective food safety management therefore requires systematic monitoring of animal feed and fodder for hazardous levels of As, Pb, Hg, and Cd [16,25]. In Bangladesh, accelerating industrialization and agricultural intensification have heightened environmental pollution pressures; however, the status of HMs in the bovine food chain—from cattle feed and fodder through to raw milk and beef—remains poorly characterized. Critically, no comparative data exist for industrial versus non-industrial zones, and the extent of bioaccumulation in cattle tissues and associated consumer health risks is largely unexplored. To address these knowledge gaps, the present study: (i) quantified concentrations of six HMs (Cd, Cr, Hg, Pb, As, and Cu) in cattle feed, fodder, raw milk, and beef from industrial and non-industrial zones; (ii) evaluated spatial heterogeneity in contamination across eight sampling sites; (iii) modelled the probabilistic trophic transfer of metals from dietary sources to animal products using Monte Carlo simulation; and (iv) assessed the associated non-carcinogenic and carcinogenic health risks for adult and child consumers.

## 2. Materials and methods

### 2.1. Study areas and study period

The study was conducted from January to April 2024 across two distinct geographical zones in Bangladesh: (i) non-industrial areas (comprising "milk pocket" and beef fattening regions) and (ii) industrial areas. In the non-industrial sector, six upazilas (sub-districts) were selected from the Sirajganj, Pabna, and Bogura districts based on livestock density data from District Livestock Offices: Shahjadpur (Site-1) and Ullapara (Site-2) in Sirajganj; Bera (Site-3) and Sathia (Site-4) in Pabna; and Sherpur (Site-5) and Shariakandi (Site-6) in Bogura. In the industrial sector, Savar (Site-7) in Dhaka and Narayanganj Sadar (Site-8) in Narayanganj were selected. Farms were targeted based on integrated milking and fodder production, while markets were chosen for high slaughter throughput.

### 2.2. Sample size and sample collection

A total of 280 samples were collected for heavy metal (HM) analysis: Fodder (n=88) including grasses, straw, and water hyacinth (48 non-industrial, 40 industrial); Cattle Feed (n=40) representing five commercial and local types per upazila; Raw Milk (n=76) collected aseptically and manually from dairy cows; and Beef (n=76) consisting of fresh samples obtained from local slaughter markets.

Samples were collected using sterile instruments and placed immediately into sterile polyethylene zip-lock bags (solid matrices) or sterile centrifuge tubes (milk). Temperature integrity during transit (4°C) was maintained using insulated containers with ice packs. Upon arrival at the Department of Pharmacology laboratory, Bangladesh Agricultural University (BAU), Mymensingh, milk and meat samples were stored at −20°C until analysis.

### 2.3. Sample preparation and digestion

Solid samples (fodder, beef, feed) were rinsed with deionized water, oven-dried at 60°C for 72 hours until reaching a constant weight, and homogenized into a fine powder. Microwave-assisted acid digestion was performed using an Ethos Easy Milestone system. Approximately 0.2 g of solid sample (or 1 mL of milk) was digested with 8 mL of 70% HNO₃ and 2 mL of 30% H₂O₂. Post-digestion, solutions were filtered (Whatman No. 41) and diluted to 50 mL with deionized water.

### 2.4. Instrumental analysis and quality control

Heavy metal concentrations (As, Cd, Cu, Pb, Hg, Cr) were determined using an Atomic Absorption Spectrophotometer (AAS; PinAAcle 900H, PerkinElmer, USA) via Graphite Furnace (GFAAS) where applicable. Analytical quality was maintained through blanks, triplicate analyses, and Certified Reference Materials (CRMs) from Inorganic Ventures (USA). The instrument was calibrated using five-point linear calibration curves established via serial dilution of 1000 mg/L stock solutions. The operating parameters for the AAS are detailed in Table 1.

**Table 1.**
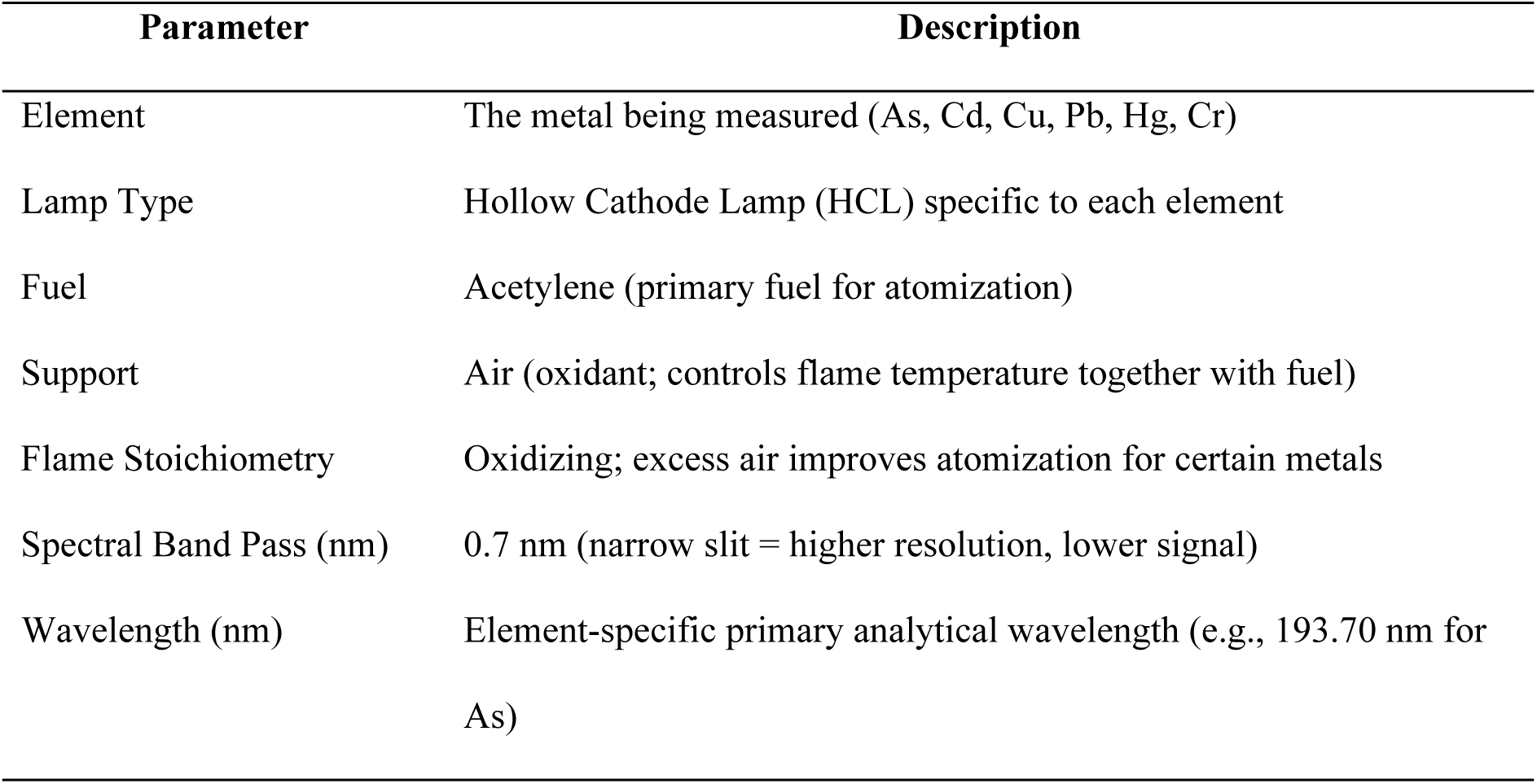
Atomic absorption spectrophotometer operating parameters for heavy metal analysis.

### 2.5. Ethical statement

The research protocol was approved by the Animal Welfare and Experimentation Ethics Committee of Bangladesh Agricultural University, Mymensingh (approval number: AWEEC/BAU/20(76); date: 14 November 2022).

### 2.6. Benchmarking against maximum permissible limits (MPL)

To assess the safety of the samples, the detected HM concentrations were compared against the Maximum Permissible Limits (MPL) established by national and international regulatory bodies (Table 2).

**Table 2.** Maximum permissible limits (MPL, mg/kg) of heavy metals in beef, cow’s milk, cattle feed, and fodder.

| Product | Cr | Cd | Pb | As | Hg | Cu |
| --- | --- | --- | --- | --- | --- | --- |
| Beef | 1.0<br>(FAO/WHO, USDA) | 0.05 (WHO, EC, CAC, EU, EFSA) | 0.1 (WHO, CAC, EU, EFSA) | 0.01<br>(WHO, USA, EFSA) | 1.0 (CAC, EU, EC) | 10.0<br>(WHO) |
| Milk | 0.025<br>(WHO) | 0.01<br>(FAO/WHO, CAC, FDA) | 0.02<br>(FAO/WHO, CAC, EU) | 0.1 [22] | 0.05 (CAC) | 0.01<br>(FAO, WHO) |
| Feed | 0.5 (FDA) | 1.0 (EU, UAE) | 5.0 (EU) | 2.0 (EU, Turkey, China) | 0.1 (EU, UAE, CN) | 35.0 (EU) |
| Fodder | 5.0<br>(FAO/WHO) | 0.5 (WHO, CERSPC) | 0.3<br>(FAO/WHO) | 4.0 (EU) | 0.5<br>(FAO/WHO) | 20.0<br>(WHO) |

*Benchmark values reflect limits most commonly used in comparable heavy-metal studies of beef, milk, feed, and fodder, including [26–31]. Not every analyte–matrix pair has a single harmonized regulatory limit; literature-based benchmark values were retained where necessary for cross-study comparison.*

### 2.7. Data analysis and risk assessment

#### 2.7.1. Transfer factor (TF) analysis

Transfer factors (TFs) were calculated to evaluate the efficiency of heavy-metal transfer from dietary sources (feed and fodder) to edible animal products (beef and milk). Mean concentrations (mg kg⁻¹, wet weight) of Cr, Cu, Cd, Pb, As, and Hg measured in cattle feed, fodder, beef, and milk were used for the analysis.

The transfer factor from a dietary source to beef was defined as the ratio of the metal concentration in beef to that in the corresponding feed or fodder, expressed as:

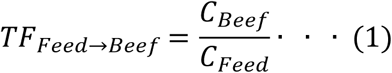

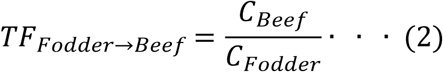

Similarly, transfer factors from dietary sources to milk were calculated using the same approach:

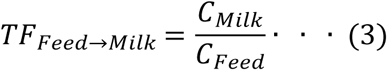

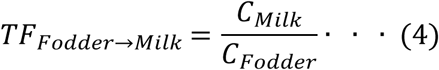

where *C*_*Beef*_, *C*_*Milk*_, *C*_*Feed*_ and *C*_*Fodder*_ represent the mean concentrations (mg kg⁻¹) of a given metal in beef, milk, feed, and fodder, respectively. In addition, feed-to-milk transfer factors were computed for descriptive purposes using the same approach, although these were not included in the main figure due to substantially lower values.

To account for uncertainty and variability in measured concentrations, Monte Carlo simulations (10,000 iterations) were performed for each metal–pathway combination. Metal concentrations in animal products and dietary sources were assumed to follow normal distributions parameterised by their observed means and standard deviations. Random samples were truncated at zero to avoid biologically implausible negative concentrations. Transfer factors were calculated iteratively as the ratio of simulated concentrations in animal products to those in the corresponding dietary source. For each pathway, the resulting TF distributions were summarised using the mean, median, and 95% uncertainty intervals (2.5th–97.5th percentiles). A TF value greater than 1 indicates potential biomagnification, whereas TF values less than 1 indicate limited transfer or effective physiological regulation [32].

#### 2.7.2. Human health risk assessment

The potential non-carcinogenic health risks associated with consumption of beef and milk contaminated with HMs were evaluated for two distinct population groups: adults and children, following the methodology established by the United States Environmental Protection Agency [33,34].

##### 2.7.2.1. Estimated daily intake (EDI)

The Estimated Daily Intake (EDI) of each heavy metal was calculated to quantify the daily exposure level per unit of body weight [35,44]:

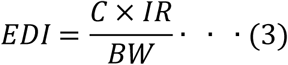

where C = mean concentration of the heavy metal in the food matrix (mg/kg for beef, mg/L for milk); IR = daily ingestion rate — based on national dietary statistics [36], the average consumption was estimated at 7.54 g/day (beef) and 27.31 g/day (milk) for adults, and 3.10 g/day (beef) and 27.31 g/day (milk) for children; and BW = average body weight, defined as 60 kg for adults and 30 kg for children [37,38].

##### 2.7.2.2. Target hazard quotient (THQ)

The non-carcinogenic risk for individual metals was assessed using the Target Hazard Quotient (THQ), the ratio of the estimated exposure to the reference dose:

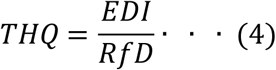

where RfD = the oral Reference Dose (mg/kg/day) sourced from the USEPA Integrated Risk Information System (IRIS): Cd (0.001), As (0.00006; updated USEPA IRIS 2025 inorganic arsenic value), Hg (0.0003), Cr (1.5), and Cu (0.04). Lead (Pb) was excluded from THQ and HI calculations because a contemporary threshold-based oral RfD is not appropriate for Pb dietary-risk screening; Pb exposure is reported as EDI only. A THQ < 1 indicates that the exposed population is unlikely to experience adverse health effects, whereas a THQ > 1 signifies a potential non-carcinogenic health risk [33,34].

#### 2.7.3. Hazard index (HI)

To evaluate the cumulative potential for adverse health effects resulting from simultaneous intake of multiple heavy metals, the Hazard Index (HI) was calculated by summing the THQ values for each specific food matrix:

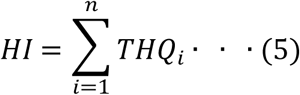

where THQi is the target hazard quotient of an individual metal. An HI value exceeding 1.0 suggests a significant risk of non-carcinogenic effects from the contaminant mixture [39].

#### 2.7.4. Lifetime carcinogenic risk (CR) from arsenic and lead

The potential lifetime carcinogenic risk (CR) of As and Pb via dietary exposure was estimated for beef and milk following the methodology recommended by the USEPA [33,40]. The EDI for each metal for adults was calculated as:

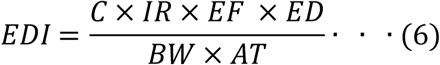

where C = metal concentration (mg/kg for meat, mg/L for milk); IR = ingestion rate (7.54 g/day for beef, 27.31 g/day for milk); EF = exposure frequency (365 days/year); ED = exposure duration (70 years); BW = average adult body weight (60 kg); and AT = averaging time (365 × 70 days) [33].

The lifetime CR for each metal was calculated as:

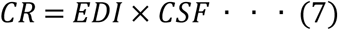

with cancer slope factors (CSF) from USEPA IRIS: 1.5 (mg/kg/day)⁻¹ for As and 0.0085 (mg/kg/day)⁻¹ for Pb. The total CR for each metal was computed as the sum of CRs from beef and milk. CR < 1×10⁻⁶ was considered negligible; values between 1×10⁻⁶ and 1×10⁻⁴ were regarded as acceptable; and values exceeding 1×10⁻⁴ indicated a potential carcinogenic concern.

#### 2.7.5. Software and supplementary file

All statistical analyses, probabilistic Monte Carlo simulations, and health risk calculations were conducted in R (Version 4.5.0) [41]. The complete annotated R code used for trophic transfer factor analysis and human health risk assessment (THQ, HI, and CR) is provided in Supplementary File 1.

## 3. Results

### 3.1. Occurrence and distribution of heavy metals in dietary sources and animal products

Table 3 presents a statistical summary of HM concentrations across all four sample matrices. Copper (Cu) was the predominant contaminant in every matrix, with mean concentrations far exceeding those of the other five metals. In beef, the descending contamination order was Cu > Pb > Cr > As > Cd > Hg. In raw milk, the hierarchy was Cu > Hg > Pb > As > Cr > Cd. Cattle feed samples similarly exhibited Cu dominance, followed by Cr > Pb > Cd > As > Hg. Fodder samples showed the same Cu-dominant pattern, with remaining metals ranked as Cr > Pb > As > Cd > Hg. Notably, Cu concentrations in beef (mean 103.9 mg/kg) and milk (mean 13.7 mg/L) substantially exceeded the maximum permissible limits (MPL) shown in Table 2.

**Table 3.**
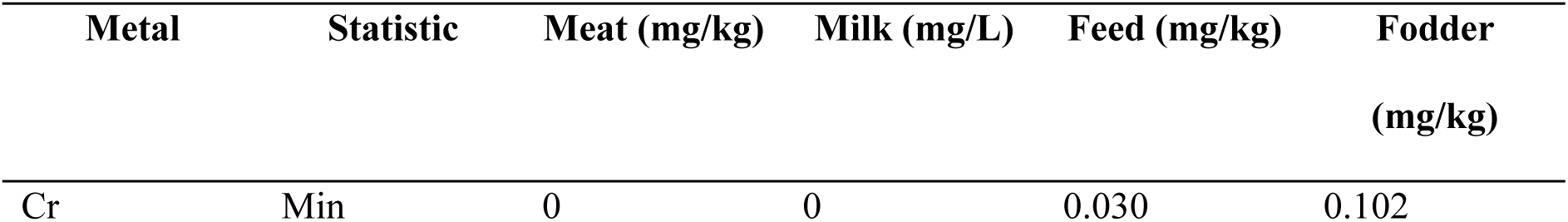
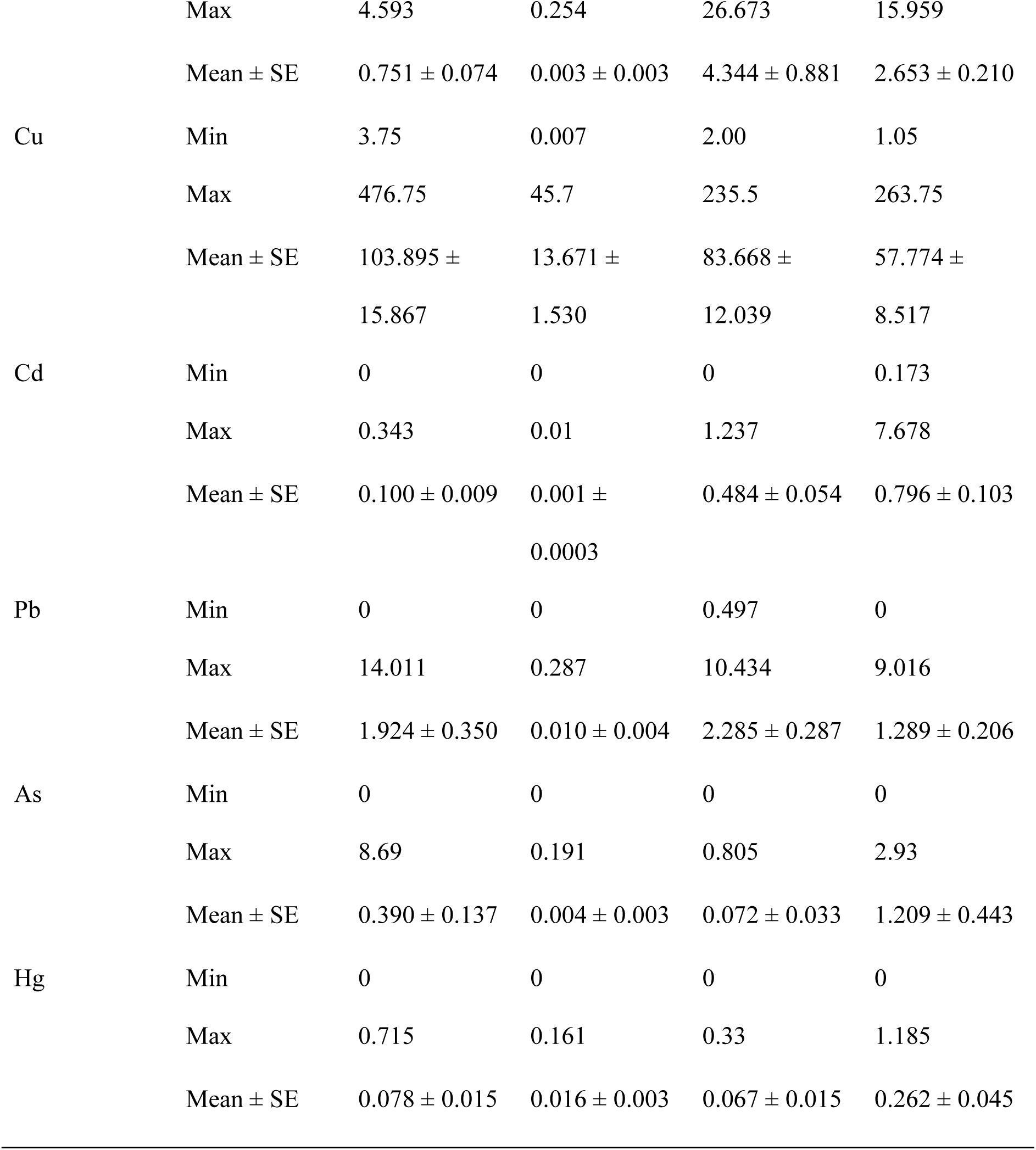
Statistical summary of heavy metal concentrations (Min, Max, Mean ± SE) in meat, milk, feed, and fodder.

### 3.2. Spatial variation of heavy metal contamination across industrial and non-industrial sites

Spatial analysis revealed marked heterogeneity in metal accumulation across the eight study sites (Supplementary Files 1–4). In industrial zones, Site-7 (Savar) and Site-8 (Narayanganj) recorded the highest mean Pb concentrations in beef (2.985 and 4.193 mg/kg, respectively), both substantially exceeding the MPL of 0.1 mg/kg. By contrast, Cr concentrations peaked in the non-industrial zone of Site-2 (Ullapara, Sirajganj; 1.257 mg/kg). Despite this spatial heterogeneity, Cu remained the most prevalent metal at all sites and across all matrices, with the highest mean concentration observed in fodder at Site-1 (77.53 mg/kg).

#### 3.2.1. Contamination profiles in beef and milk

Copper (Cu) consistently exhibited the highest residue levels in beef across all sampling sites, reaching concentrations of up to ∼185 mg/kg. In contrast, mercury (Hg) and chromium (Cr) were detected at very low or near-background levels, indicating minimal presence relative to other metals (Figure 1).

**Figure 1.**
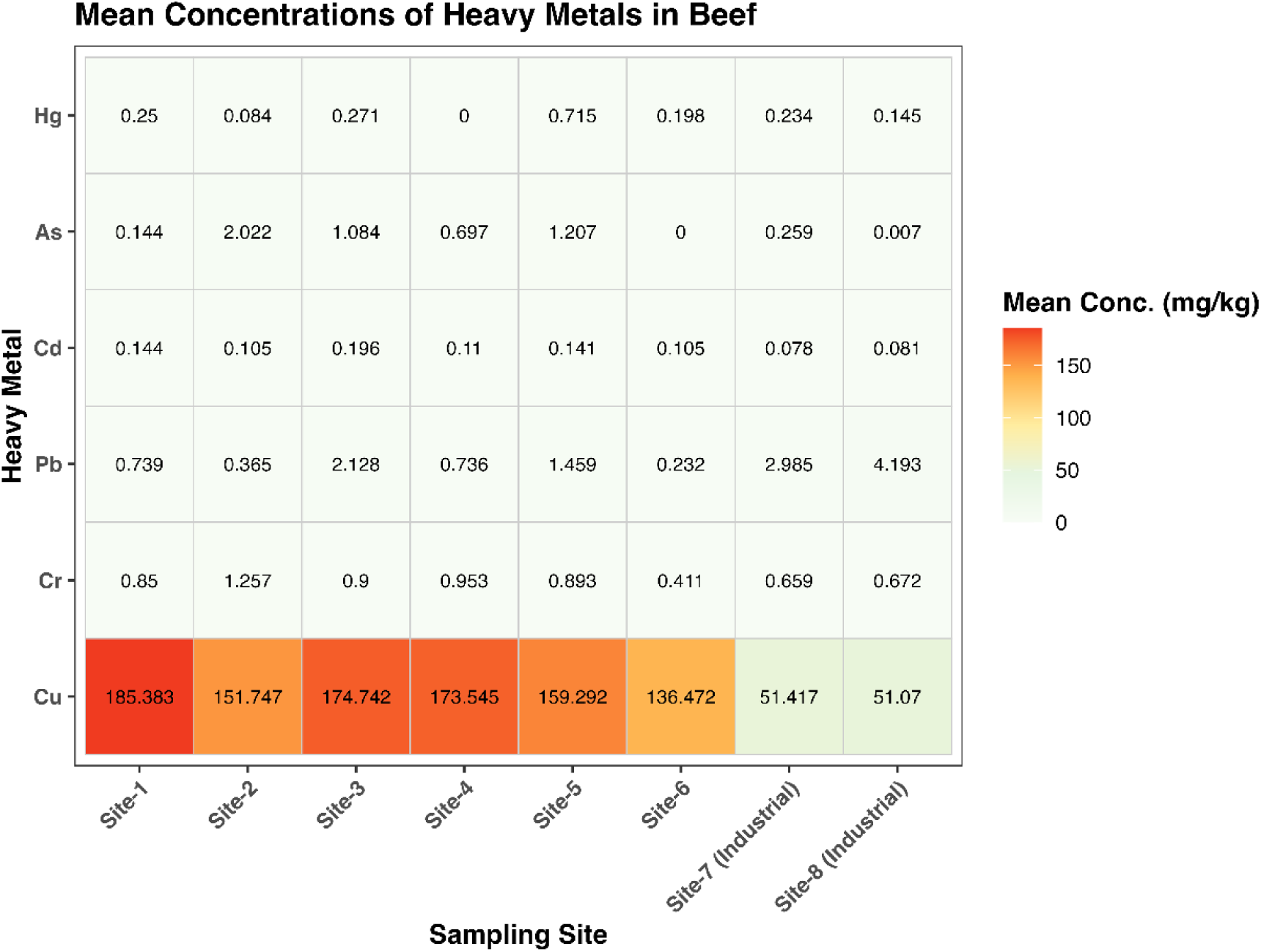
Site-specific heavy metal residues (Cr, Cu, Cd, Pb, As, and Hg) detected in beef samples (mg/kg). The heat map compares contamination patterns between locations, with darker shades representing higher concentrations of the respective metals.

A similar trend was observed in milk samples, where Cu was the most prevalent metal, with concentrations markedly higher than those of other analytes. The peak Cu concentration was recorded at Site-3 (20.694 mg/L). Conversely, Hg, Cr, and Pb were detected at low concentrations or were absent (0 mg/L) at several sites (Figure 2).

**Figure 2.**
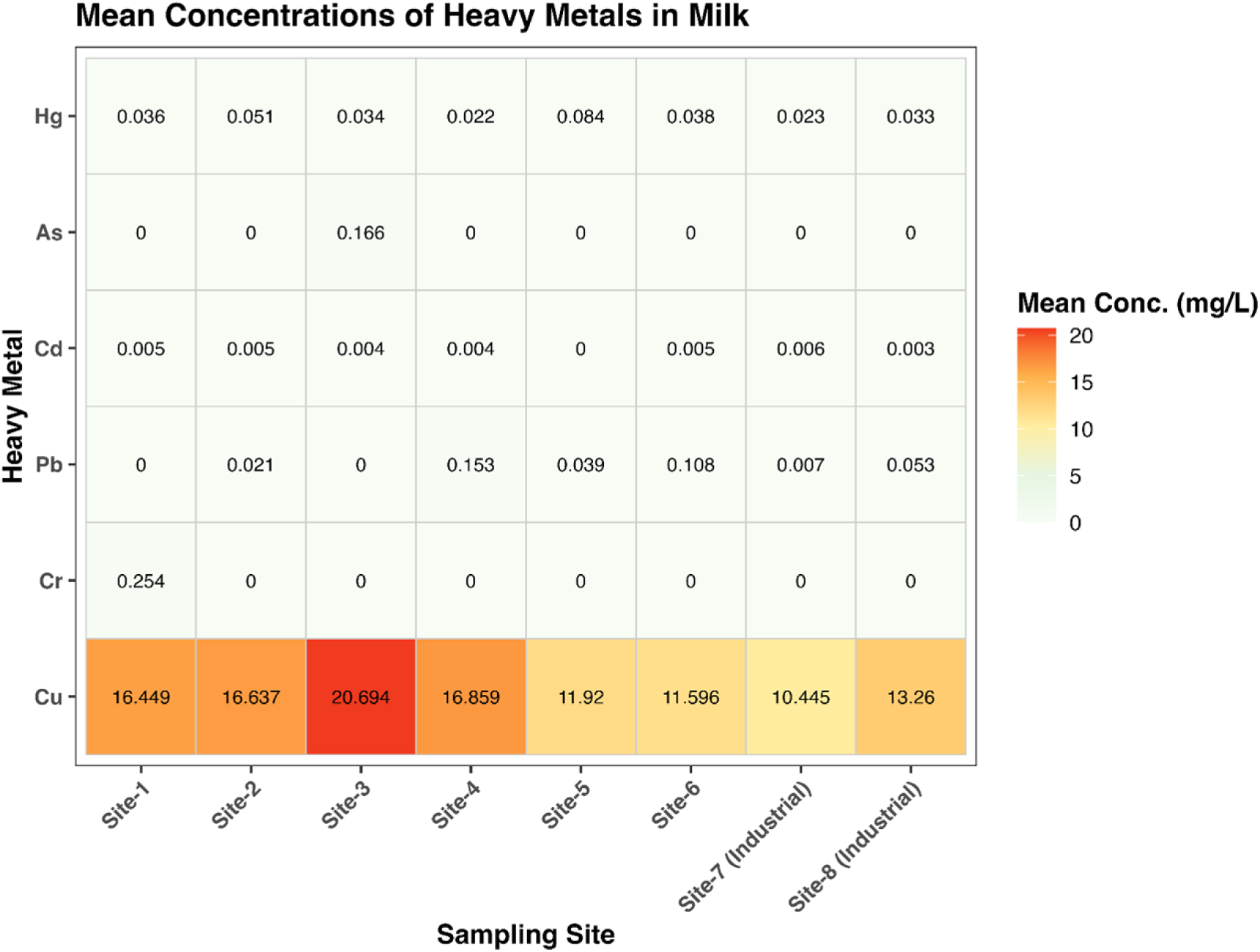
Site-specific heavy metal residues (Cr, Cu, Cd, Pb, As, and Hg) detected in milk samples (mg/L). The heat map compares contamination patterns between locations, with darker shades representing higher concentrations of the respective metals.

#### 3.2.2. Heavy metal burden in cattle feed and fodder

Copper (Cu) was the predominant contaminant in cattle feed, displaying significantly higher concentrations than all other metals. A distinct contamination hotspot was observed at Site-5, where the Cu concentration peaked at 158 mg/kg. The remaining metals, particularly Cr and Pb, showed substantially lower levels, while As, Cd, and Hg were minimal, confirming Cu as the primary driver of heavy metal burden in feed (Figure 3).

**Figure 3.**
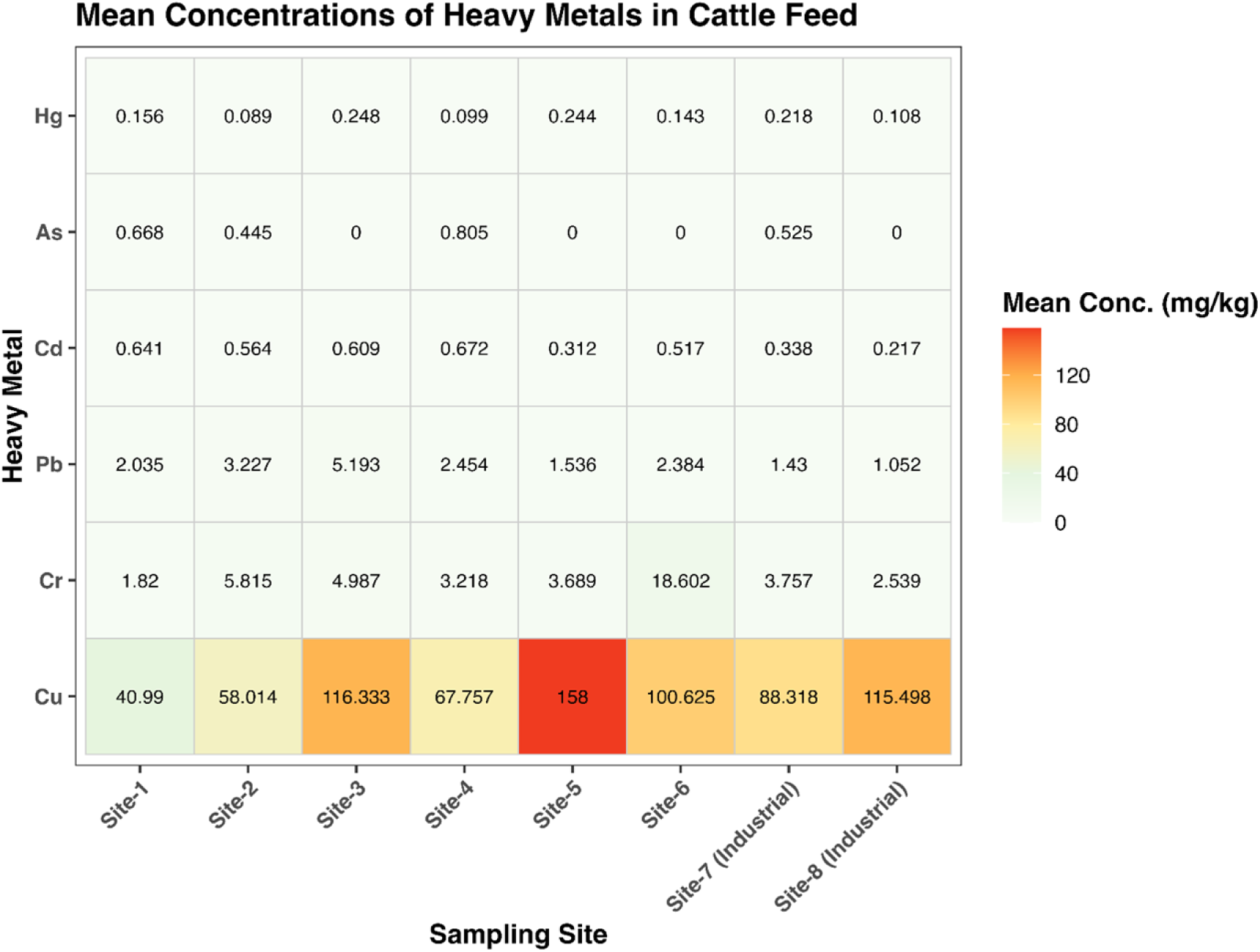
Spatial distribution of heavy metals in cattle feed across different sampling sites. The heat map displays the relative concentration profiles of Cd, Cr, Hg, As, Pb, and Cu, with colour intensity indicating higher contamination levels.

Significant spatial variability was also observed in fodder samples (Figure 4). Cu was identified as the dominant pollutant, with Sites-1, 2, and 4 acting as distinct hotspots exhibiting exceptionally high concentrations (>74.05 mg/kg). In contrast, levels of the other five metals (Cd, Pb, As, Hg, and Cr) remained substantially lower and relatively consistent across all sites.

**Figure 4.**
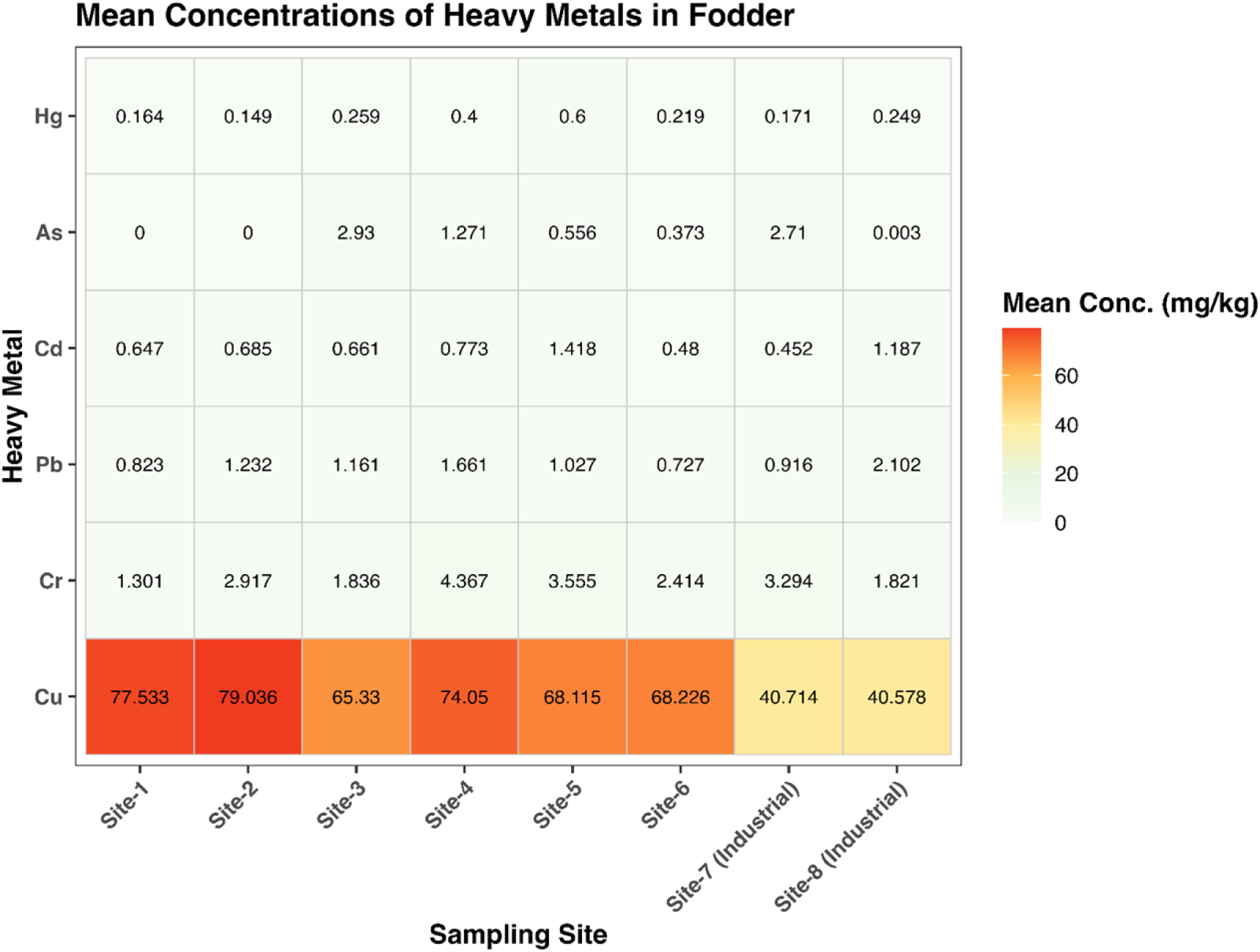
Spatial distribution of heavy metals in fodder samples across different sampling sites. The heat map displays the relative concentration profiles of Cd, Cr, Hg, As, Pb, and Cu (mg/kg), with colour intensity indicating higher contamination levels.

### 3.3. Probabilistic trophic transfer of heavy metals from dietary inputs to beef and milk

Figure 5 illustrates the simulated distributions of Transfer Factors (TF) for Cr, Cu, Cd, Pb, As, and Hg across feed-to-beef and fodder-to-beef pathways, derived from Monte Carlo simulation. The violin plots reveal substantial variability in transfer efficiency among metals and between pathways. Copper (Cu) demonstrated the highest accumulation potential, with a median transfer efficiency (TF) exceeding 1.0. In contrast, Cr and As showed lower transfer efficiencies, with most distributions falling below 1.0. Overall, feed proved to be a more efficient vector for heavy metal translocation into beef compared to fodder.

**Figure 5.**
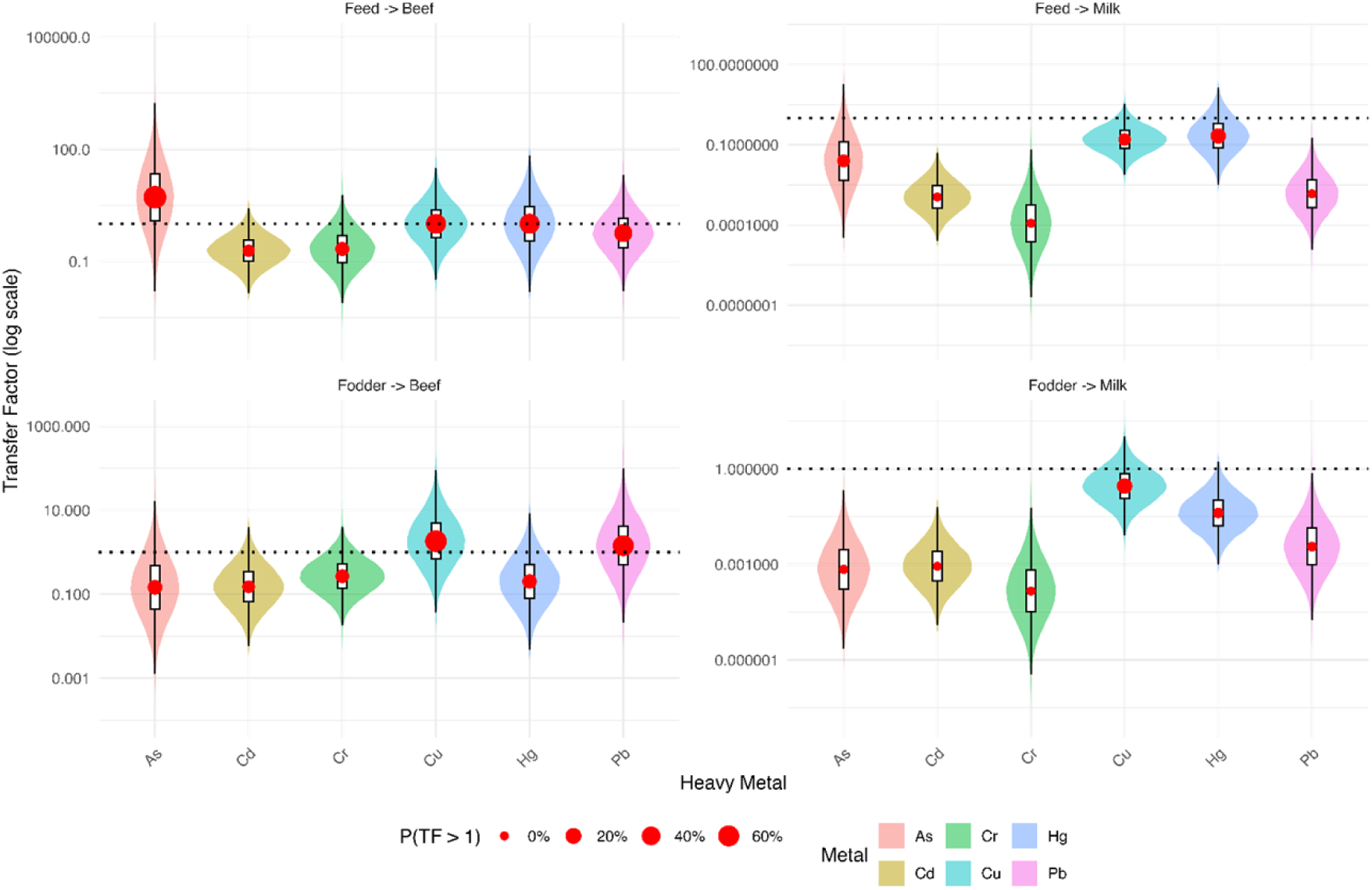
Probabilistic distributions of heavy metal transfer factors (TF) from feed and fodder into beef and milk, derived from Monte Carlo simulation (10,000 iterations).

### 3.4. Non-carcinogenic public health risk

The non-carcinogenic health risks associated with beef and milk consumption were evaluated using the Target Hazard Quotient (THQ) model (Figure 6). Risk estimates are presented separately for adults and children, with box plots differentiating between beef (brown) and milk (blue) exposure pathways. Copper (Cu) was identified as the predominant hazard, being the only metal to consistently exceed the safety threshold (THQ = 1). While Cu indicates a moderate non-carcinogenic risk for adults, THQ values are significantly higher for children, denoting a substantially elevated risk of overexposure through these food sources.

**Figure 6.**
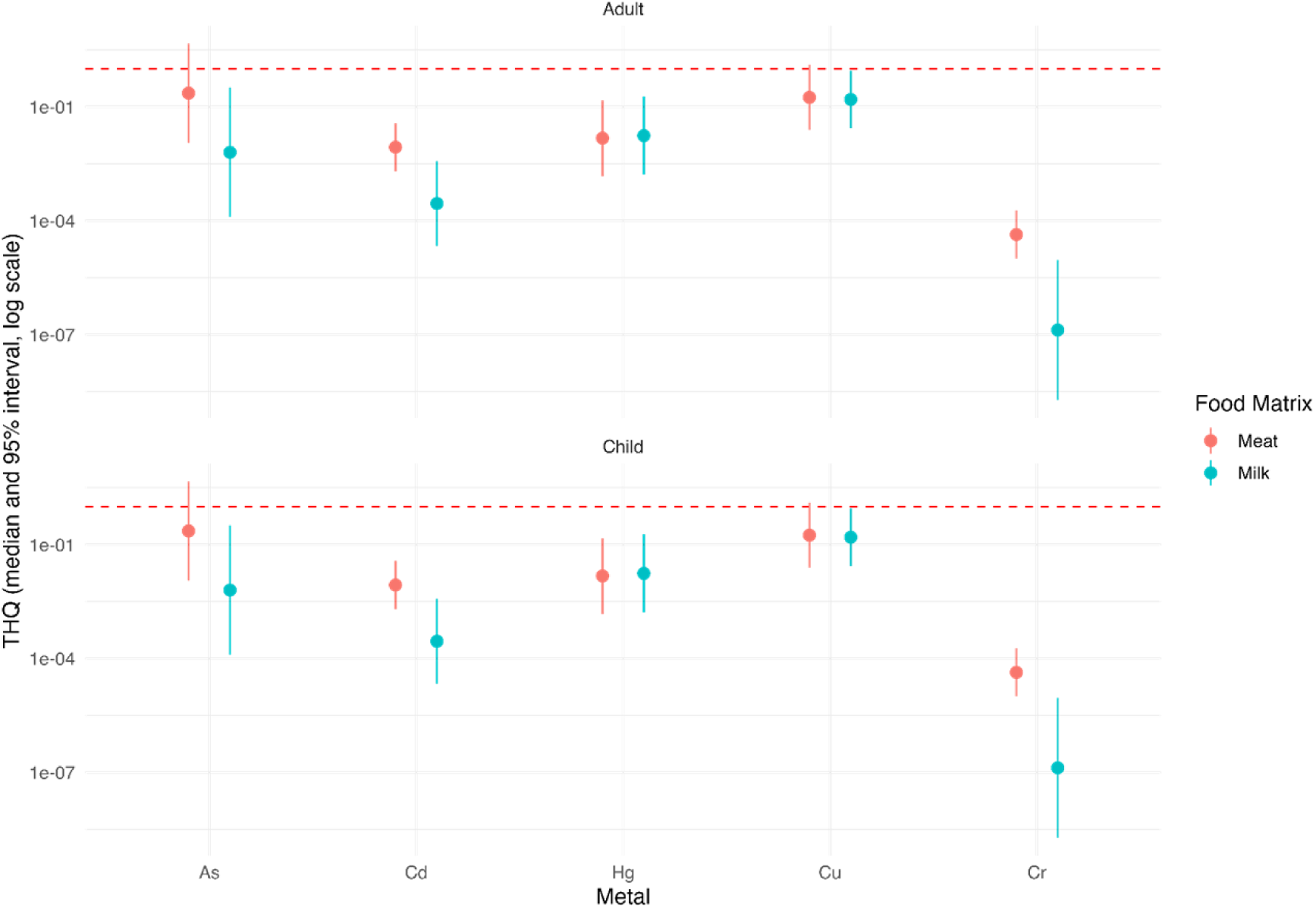
Box plots illustrating the non-carcinogenic health risks, expressed as the Target Hazard Quotient (THQ), for adults and children consuming beef and milk contaminated with heavy metals (As, Cd, Cr, Cu, Hg, and Pb). The y-axis is presented on a log₁₀ scale, and the red dashed line indicates the critical safety threshold (THQ = 1). Brown boxes represent beef; blue boxes represent milk.

Figure 7 presents the cumulative non-carcinogenic risk, expressed as the Hazard Index (HI), for both adults and children. The box plots compare the combined contributions of all analysed heavy metals across the two food matrices. Overall, HI values were consistently higher for children than for adults, indicating greater vulnerability to cumulative toxicological exposure. All estimates were evaluated relative to the safety threshold of HI = 1.

**Figure 7.**
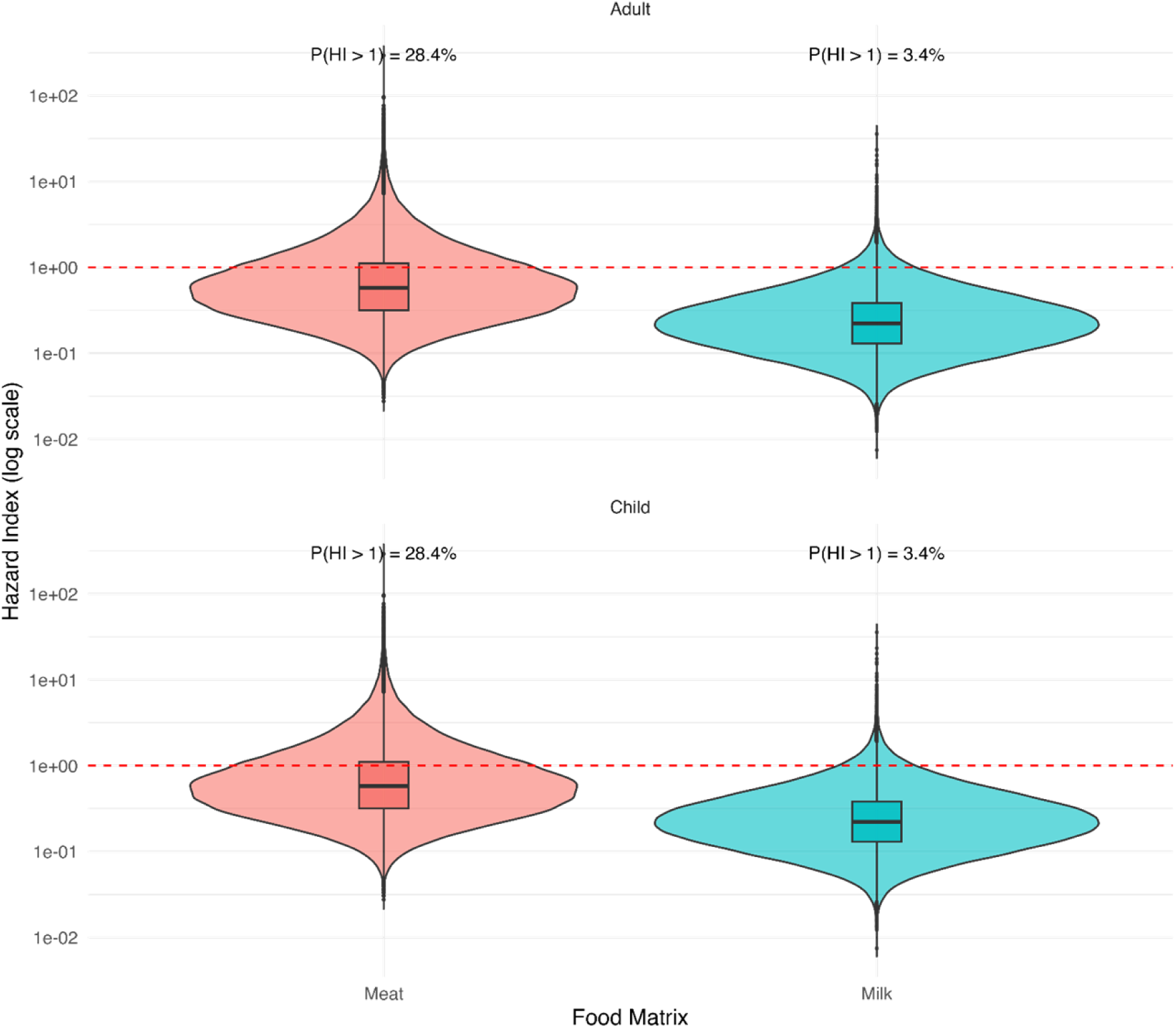
Comparison of the cumulative Hazard Index (HI) for adults and children consuming beef and milk. Box plots represent the combined non-carcinogenic risk of all target heavy metals on a log₁₀ scale. The red dashed line indicates the safety threshold (HI = 1), above which significant health concerns exist.

### 3.5. Lifetime carcinogenic risk

The lifetime carcinogenic risks (CR) for adults resulting from dietary exposure to arsenic (As) through beef and milk consumption are presented in Figure 8. The assessment reveals that arsenic poses the more significant threat, with risk distributions for both beef and milk clustering near the upper threshold of 10⁻⁴. In contrast, lead-associated risks were generally lower, with beef showing moderate proximity to the concern threshold and milk predominantly falling within the negligible risk range.

**Figure 8.**
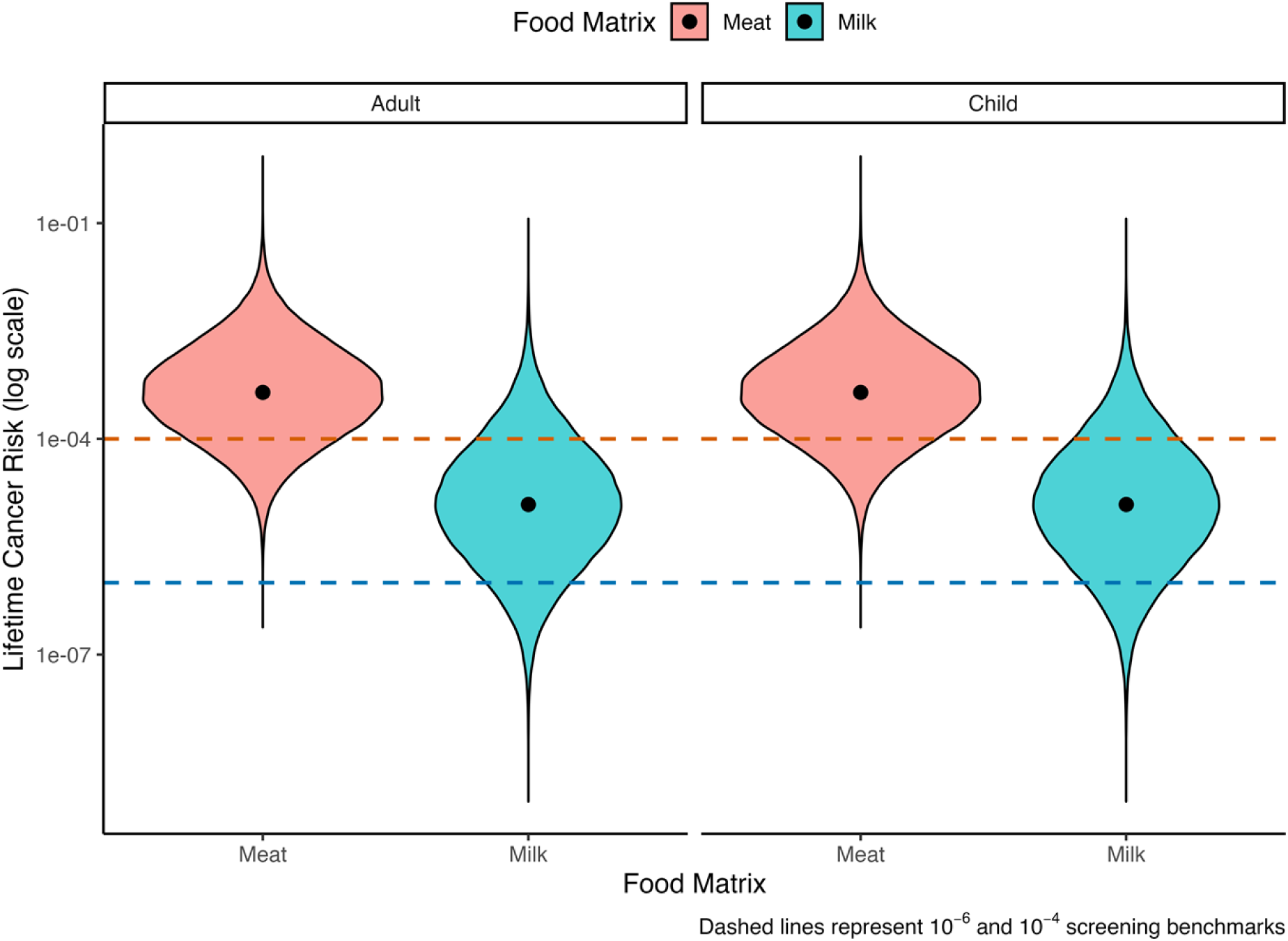
Lifetime carcinogenic risk (CR) for adults resulting from dietary exposure to arsenic through contaminated beef and milk consumption.

## 4. Discussion

This study provides the first comprehensive assessment of HM contamination, spatial distribution, trophic transfer dynamics, and associated human health risks within Bangladesh’s bovine food chain. Copper (Cu) emerged as the single dominant contaminant across all four matrices, with mean concentrations in beef, milk, feed, and fodder far exceeding those of the five co-investigated metals. Lead (Pb) and chromium (Cr) displayed contrasting spatial patterns—Pb concentrations were highest in industrial zones, while Cr peaked in a non-industrial zone—indicating that contamination source profiles differ markedly between land-use contexts. Probabilistic transfer modelling identified commercial feed as the primary vector for HM translocation into beef. Risk assessment confirmed Cu as the principal driver of non-carcinogenic risk, especially for children, while arsenic (As) constitutes the dominant carcinogenic hazard.

Copper emerged as the single most pervasive contaminant across all four matrices in this study, with mean concentrations in beef (103.9 mg/kg), milk (13.7 mg/L), feed (83.7 mg/kg), and fodder (57.8 mg/kg) substantially exceeding those of the five other target metals. Several interconnected sources likely explain this pattern. Copper sulfate is routinely used in commercial livestock feed as a growth promoter and antifungal agent, and the widespread use of copper-containing fungicides on fodder crops in Bangladeshi agriculture provides a continuous plant-based entry pathway. Industrial effluent releases in peri-urban zones may further elevate ambient soil-copper levels accessible to grazing animals. This finding is broadly consistent with reports from other South Asian settings where Cu frequently dominates the heavy-metal profile of cattle feed and edible products [5,16,42,43]. Although Cu is an essential trace element, chronic dietary intake above reference values is associated with hepatotoxicity and oxidative stress, underscoring the public health relevance of the high concentrations observed here.

Spatial analysis revealed marked heterogeneity in the distribution of Pb and Cr that cannot be explained by feed composition alone. Lead concentrations peaked in beef sampled from Sites 7 and 8 (Savar and Narayanganj), two of Bangladesh’s most densely industrialised districts, where tanneries, battery recycling facilities, and heavy vehicle traffic contribute to elevated atmospheric and soil-lead burdens. In contrast, Cr was highest in the non-industrial milk-pocket area of Site 2 (Ullapara, Sirajganj), a pattern consistent with the use of Cr-contaminated irrigation water or locally sourced fertilisers rather than with direct industrial fallout. These contrasting spatial signatures argue for source-specific regulatory responses: Pb monitoring should prioritise livestock operations adjacent to industrial zones, whereas Cr surveillance may need to extend to agricultural areas where contaminated water resources are used for fodder irrigation.

The probabilistic transfer-factor modelling confirms that feed is a more efficient vector for heavy-metal translocation into beef than fodder, with Cu and Hg showing the highest and most consistent median transfer efficiencies. This distinction has direct practical implications: commercial feed formulation and raw ingredient sourcing represent concentrated intervention points where regulatory limits and supplier audits could substantially reduce the metal burden entering the food chain. Feed ingredients such as fish meal, mineral premixes, and grain by-products are known to vary widely in heavy-metal content depending on geographic origin and processing practices; mandatory pre-market testing of these ingredients would therefore be an efficient upstream control measure. The comparatively lower transfer factors for fodder into milk are consistent with physiological dilution in the mammary gland and selective biliary excretion, suggesting that direct feed is the more urgent regulatory target for bovine products in Bangladesh.

Non-carcinogenic risk assessment using corrected dietary intake rates based on Bangladesh national survey data [36,37] showed that median Hazard Index values remained below 1 for all consumer scenarios. However, the upper tail of the simulated distribution exceeded unity for adult beef consumers and in the child milk sensitivity scenario, driven predominantly by Cu. Children face a disproportionately elevated relative exposure because their lower body weight amplifies dose per kilogram, and their developing organ systems are more susceptible to metal toxicity. These findings reinforce the need for dietary guidance that specifically addresses consumption of beef and milk by children in high-exposure settings [45]. Pb is not included in the hazard index because no scientifically defensible threshold-based oral reference dose exists for Pb in contemporary risk-screening frameworks; its estimated daily intake is reported separately as a qualitative food-safety concern.

Arsenic represents the dominant carcinogenic concern in this dataset. Upper-bound lifetime cancer-risk estimates for arsenic from beef consumption approach the USEPA benchmark of 10⁻⁴, a threshold at which regulatory action is typically triggered. The high arsenic loading in beef relative to milk is consistent with differential bioaccumulation in muscle tissue and longer biological residence times compared with secretion into milk. It is important to note that this is an upper-bound screening calculation assuming total arsenic behaves as inorganic arsenic; speciation data are required to refine this estimate, because organic arsenicals common in animal feed carry lower carcinogenic potency than inorganic As(III) and As(V). Notwithstanding this uncertainty, the proximity of risk estimates to the 10⁻⁴ threshold is sufficient to classify arsenic as a priority contaminant warranting confirmation through targeted speciation analysis and, if confirmed, immediate regulatory intervention on arsenic-containing feed additives.

## 5. Conclusions

This study demonstrates that HM contamination of the bovine food chain in Bangladesh is driven by multiple, spatially heterogeneous sources rather than a simple industrial versus non-industrial divide. Copper exhibited the highest burden across all matrices (beef, milk, feed, and fodder), while mercury was consistently the lowest. Probabilistic transfer-factor modelling confirmed that commercial feed is a more efficient vector for HM translocation into beef than fodder, underscoring the importance of upstream feed ingredient quality control. Health risk assessment identified Cu as the primary driver of non-carcinogenic risk, with THQ values exceeding the safety threshold (>1) for children, and arsenic as the principal carcinogenic hazard, with upper-bound lifetime cancer risk estimates approaching the USEPA benchmark of 10⁻⁴. Although median Hazard Index values remained below unity, upper-tail risk estimates and the arsenic carcinogenic screen collectively justify continued and expanded surveillance. Future work should prioritise metal speciation analysis (particularly for arsenic), longitudinal feed–water–fodder tracing, and raw sample-level probabilistic modelling to refine exposure estimates. These findings provide critical baseline evidence to inform regulatory limits for HMs in livestock feed and fodder in Bangladesh and call for immediate policy action to safeguard public health, with particular attention to protecting children from Cu and As dietary exposure.

## Author contributions

K.R. (Kazi Rafiq) and S.M.I. (Shah Md. Iqbal) contributed equally to this work. S.M.I.: Writing – original draft. K.R., A.K.M.A.R. (A K M Anisur Rahman), and M.S.I. (Md. Shafiqul Islam): Conceptualization, Supervision. K.R., A.K.M.A.R., and M.T.H. (Muhammad Tofazzal Hossain): Writing – review & editing. S.M.I. and F.S.S. (Fardina Sultana Sumi): Data curation. S.M.I. and M.R.H. (Md. Rakib Hasan): Formal analysis. M.R.H., A.B.Z. (Anan Binte Zaman), and S.M.I.: Visualization. All authors have read and approved the final manuscript.

## Acknowledgements

The authors wish to convey their profound gratitude to the Bangladesh Agricultural University Research System (BAURES), Mymensingh, for project management support and partial financial assistance toward publication expenses. The authors also extend sincere thanks to the PMHCL, BAU, Mymensingh, and the Central Laboratory of Fisheries Department, PSTU, Patuakhali, for their collaboration in sample preparation, digestion, and analyses. Additionally, the authors acknowledge the district livestock officers, upazila livestock officers, veterinary surgeons, and livestock extension officers for their support in the study areas, and all respondents—especially the large-animal farmers—for their generous cooperation during sample collection.

## Data availability statement

All data generated or analysed during this study are included in the manuscript’s Tables and Figures. The complete annotated R code used for trophic transfer factor analysis and human health risk assessment (THQ, HI, and CR) is provided in Supplementary File 1.

